# SAI: Fast and automated quantification of stomatal parameters on microscope images

**DOI:** 10.1101/2022.02.07.479482

**Authors:** Na Sai, James Paul Bockman, Hao Chen, Nathan Watson-Haigh, Bo Xu, Xueying Feng, Adriane Piechatzek, Chunhua Shen, Matthew Gilliham

## Abstract

Using microscopy to investigate stomatal behaviour is a common technique in plant physiology research. Manual inspection and measurement of stomatal features is a low throughput process in terms of time and human effort, which relies on expert knowledge to identify and measure stomata accurately. This process represents a significant bottleneck in research pipelines, adding significant researcher time to any project that requires it. To alleviate this, we introduce StomaAI (SAI): a reliable and user-friendly tool that measures stomata of the model plant Arabidopsis (dicot) and the crop plant barley (monocot grass) via the application of deep computer vision. We evaluated the reliability of predicted measurements: SAI is capable of producing measurements consistent with human experts and successfully reproduced conclusions of published datasets. Hence, SAI boosts the number of images that biologists can evaluate in a fraction of the time so is capable of obtaining more accurate and representative results.

## Introduction

Stomata, derived from the Greek word *mouth*, are small pores penetrating the epidermal surface of plant aerial organs. In monocot grasses, such as barley or maize, the stomatal apparatus includes a pair of subsidiary cells flanking the dumbbell-shaped guard cells surrounding the stomatal pore^1,2^. Dicot plants, instead, have a pair of kidney-shape guard cells surrounding each stomatal pore. Stomatal pores play a critical role in plant physiology by limiting the diffusion of carbon dioxide (CO_2_) into leaves, which directly impacts the rate of photosynthesis. Photosynthesis produces the carbohydrates, adenosine triphosphate (ATP), and nicotinamide adenosine phosphate (NADPH) required for plant metabolic functions, growth and development; releasing oxygen as a by-product. At the same time, water vapour released via stomatal pores enables water transport through plants^3,4^. Some plants survive during excessive heat by keeping stomata open, cooling leaves through the evaporation of water. Conversely, stomata are closed during droughts to prevent water loss^4^. Stomata also respond to diel cycles, such as light and dark, and a multitude of other signals to optimize CO_2_ gain and water loss^5,6^. As a consequence, stomatal aperture regulation during daily light and dark cycles, or in response to environmental stresses, directly impacts plant growth, development and survival^6–8^.

Due to the important role that stomata play, investigating stomatal regulation has become a common task for biologists studying plant signalling pathways and stress perception^6,9,10^. To study stomata traits (i.e., size or density) researchers commonly use microscopy^11–13^. This method of examining stomatal behavior, although commonplace, is not straightforward. Morphological differences in stomata of different species (Figure 1) and variable image quality make accurate stomatal measurement a task that requires experience and training. Traditionally, stomatal measurement requires manual inspection of each image to identify and measure relevant features (i.e. stomatal pore area and aperture). Hundreds of images need to be analyzed this way to gain sufficient statistical power to support a biological conclusion; a time-consuming and laborious process. Although manual measurement can be aided by image processing software such as Fuji-ImageJ^14^, manually tuned parameters are required to produce acceptable performance^15^. An automated stomatal measurement system is thus highly desirable and will accelerate plant physiology research.

**Fig. 1:**
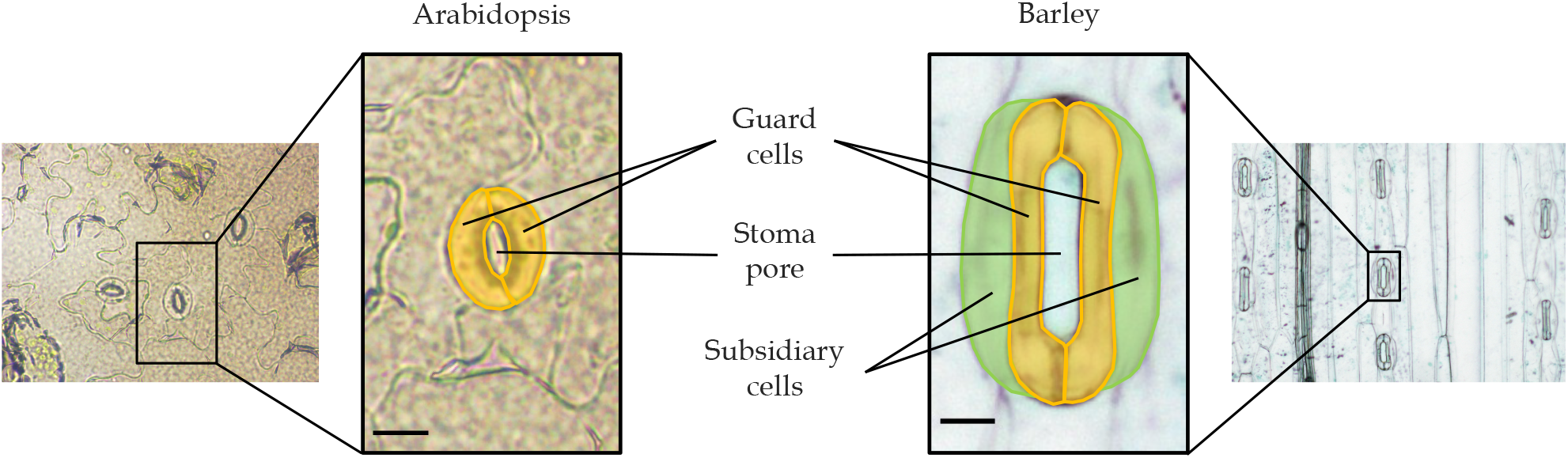
The stomata of *Arabidopsis thaliana* and barley (*Hordeum vulgare*). Components of Arabidopsis and barley stoma are highlighted and labelled; the scale bar equates to 10 μm for both Arabidopsis and barley.

Microscopy imaging presents a uniquely controlled environment for the application of modern computer vision techniques. Images can be captured in high-resolution via calibrated optics, reducing systematic noise, and plant anatomy enforces regularity in pattern, appearance and orientation (in monocot grasses). These factors remove several of the common *Achilles’ Heels* of applied vision systems. Previous attempts have been made to quantify stomatal attributes using traditional computer vision techniques to predict stomatal density, width and area^16–21^. Although these methods demonstrate efficacy on their respective tasks, they rely on handcrafted and/or multi-stage processes. The use of Convolutional Neural Networks (CNNs) to detect stomatal attributes has recently increased in popularity^22–29^. CNNs enable a series of pertinent operations to be learnt from examples - acting as a data driven approximation of a sequence of computer vision operations.

More recently, Mask Regions with Convolutional Neural Network features (Mask R-CNN) has been used to perform identification and localisation of stomata. This involves the entire stomatal complex being detected, encircled by a polygon with its orientation and stomatal complex area captured, inferring axis length^30^ or stomata density^31^. The algorithms were successfully used across different species with varying image quality^30^. Of the techniques surveyed, many studies only estimate stomatal counts for use in density calculations,^17,19,21,25,26,29,31^ with Fetter et al. (2019)^26^ providing a user-friendly online application named “Stomata Counter”. Fewer studies are focused on stomatal pore measurements, with methods that are semi-automated requiring handcrafted feature extractors or manual post-processing following model inference^18,22,24,27,28^. Ellipse fitting is the common solution used among these studies for estimating the pore area, width and length through calculating the fitted ellipse’s area, minor-axis and major-axis^22,24,27,28,32^. However, the fitting method is restricted to stomata with an ovular-shaped pore (e.g. Arabidopsis stomata), and other shapes of the stomatal pore (e.g. barley) cannot be represented correctly with an ellipse and result in under or over estimation of pore features (Figure 1). Besides, none of the above studies offers a usable automated stomatal pore measurement tool available for use.

Here, we present StomaAI (SAI) as an accessible automated tool that allows stomatal pore measurement of microscope images. The precise stomatal pore feature measurement is the core novelty of StomaAI (SAI), measuring pore area, length, width (i.e. aperture), and width/length ratio. We demonstrate that measurements obtained using SAI are comparable to those taken by human experts, providing assurance of prediction reliability. This key comparison is not provided by contemporary studies that use traditional computer vision evaluation criteria such as F1 score or average precision (AP) to evaluate machine performance. Due to differences in stomata morphology, SAI includes two class-specific models: a dicot model trained with Arabidopsis data and a monocot cereal model trained with barley data. We demonstrate that with approximately 150 annotated images containing about 1700 stomata, SAI can be trained to measure pores of two different plant species. The online demonstrator software where model inference can be viewed is hosted at https://sai.aiml.team. To use SAI to measure user acquired samples, we provide a local version that can be accessed via https://github.com/xdynames/sai-app.

## Results

### SAI achieves *human-level* performance

Beyond assessing performance using traditional metrics, we show that SAI produces measurements that are equivalent to *human-level* performance. To compare independent human operators (multiple plant physiology researchers) with SAI we applied an Average-Human/Machine Test. To reduce rater’s bias, the average measurements taken by 4 human experts were used to provide a *human-level* reference. The concordance correlation coefficient (CCC; ranging from −1 to 1) was used to evaluate the agreement between different human measurements, the *human-level* reference and SAI^33^. Stomatal width, length, area and width/length ratio were measured by SAI (Figure 2, Appendix Figure A1) and human experts (Appendix Figure A2). Width measurements obtained from SAI, when plotted against reference measurements, generally align with *y = x*; indicating that width measurements are consistent with the reference (Figure 2). Incorrectly classified samples (i.e. where the predicted opening status disagree with the reference) can be identified as those points along the *x* or *y* axes. SAI achieves a CCC of 0.891 and 0.984 for Arabidopsis and barley respectively. Considering that any Arabidopsis open stomata with a stomatal width of less than 1 μm will have a minimal impact on transpiration, the Arabidopsis stomatal width achieves a CCC at 0.916 when excluding stomata that have a width of less than 1 μm *human-level* reference. Human experts show an average CCCs of 0.9449 on Arabidopsis samples and 0.9853 when measuring barley (Appendix Figure A2). Measurements performed on barley samples exhibited improved correspondence with the reference in all cases.

**Fig. 2:**
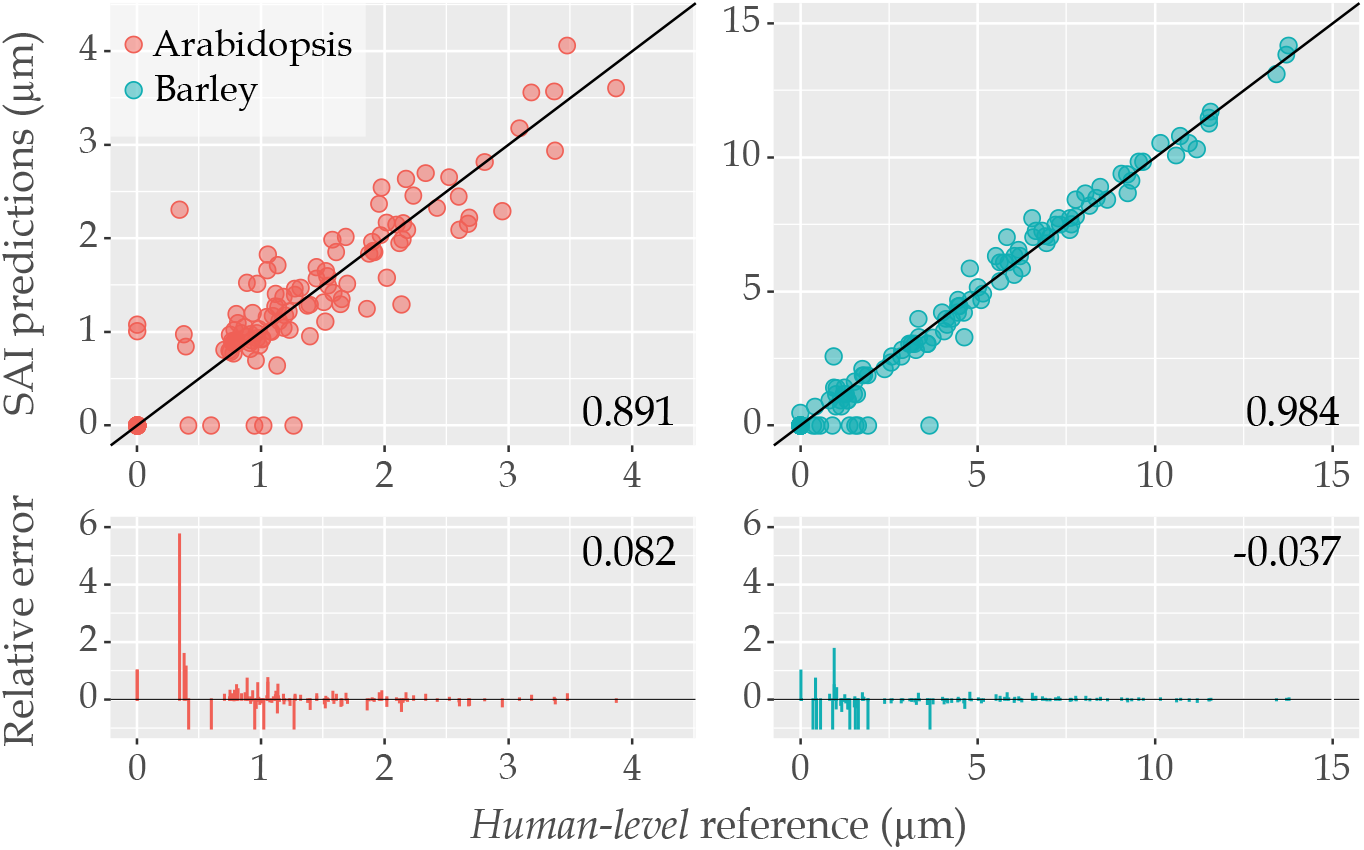
SAI prediction vs average *human-level* reference set in Arabidopsis and barley stomatal width (μm). Stomata morphology measurements from 4 human experts were collected and an average width of each stoma were calculated as the *human-level* reference. In upper panel, SAI predictions were compared against the reference and the concordance correlation coefficient (CCC, ranging from −1 to 1) was displayed as the determination of the accuracy performance. The black diagonal line represents *y = x* and CCC is a measure of dispersion for the points from that diagonal line. The corresponding relative error (RE) to *human-level* reference was presented at lower panel with mean RE presented. Data points are color coded by plant species. (Arabidopsis: *N >* 120, barley: *N >* 160)

Relative errors (RE) in measurements from SAI are distributed in a similar pattern to those from human measurements (Appendix Figure A2). Judging aperture extent in stomata that are almost closed is more difficult than when they are open. This creates a skew in the RE histogram where errors are more frequently observed in small measurements. Estimation of stomatal length were not affected by stomatal opening status, so REs are evenly spread between under and over estimation. The mean stomatal width, length, area and width/length ratio are calculated for each source of measurements and compared using one-way ANOVA with Tukey HSD (Appendix Table A1). This comparison aims to test whether measurement sources exhibit a statistically significant difference. No such significance was found in measured stomatal features across both Arabidopsis and barley samples when SAI was compared to the *human-level* reference measurements (Appendix Figure A3). Additionally, SAI exhibits no significant difference from individual human expert measurements, except in the case of expert 2’s length measurements for Arabidopsis. Interestingly, human expert 2’s Arabidopsis measurements are significantly smaller in area, length and width to both human expert 1 and 3. In all cases, expert 2 tends to measure stomata more conservatively compared to others.

### SAI produces consistent replication of human processed datasets

SAI was used to measure two sets of published physiological experiments. The original images from Xu et al. (2021)^11^ Supplementary Figure 3b & 5g were processed with SAI. Traditionally, researcher’s will exercise their discretion by consciously measuring only stomata they deem as mature. SAI measures indiscriminately. However, we were able to emulate this practice via filtering of detections based on their estimated length. To exclude immature stomata, we eliminate detections of Arabidopsis stomata with lengths shorter than 2 μm and barley stomata shorter than 16 μm in length. SAI and the original manual measurements were compared using ANOVA with Tukey HSD, as in Xu et al. (2021)^11^ (Figure 3). Scientific conclusions drawn from the statistical tests were consistent with those of Xu et al. (2021)^11^. Arabidopsis stomata are closed in response to 25 μM ABA in the presence and absence of 2 mM GABA, and light induced barley stomatal opening was inhibited by the presence of 1 mM GABA.

**Fig. 3:**
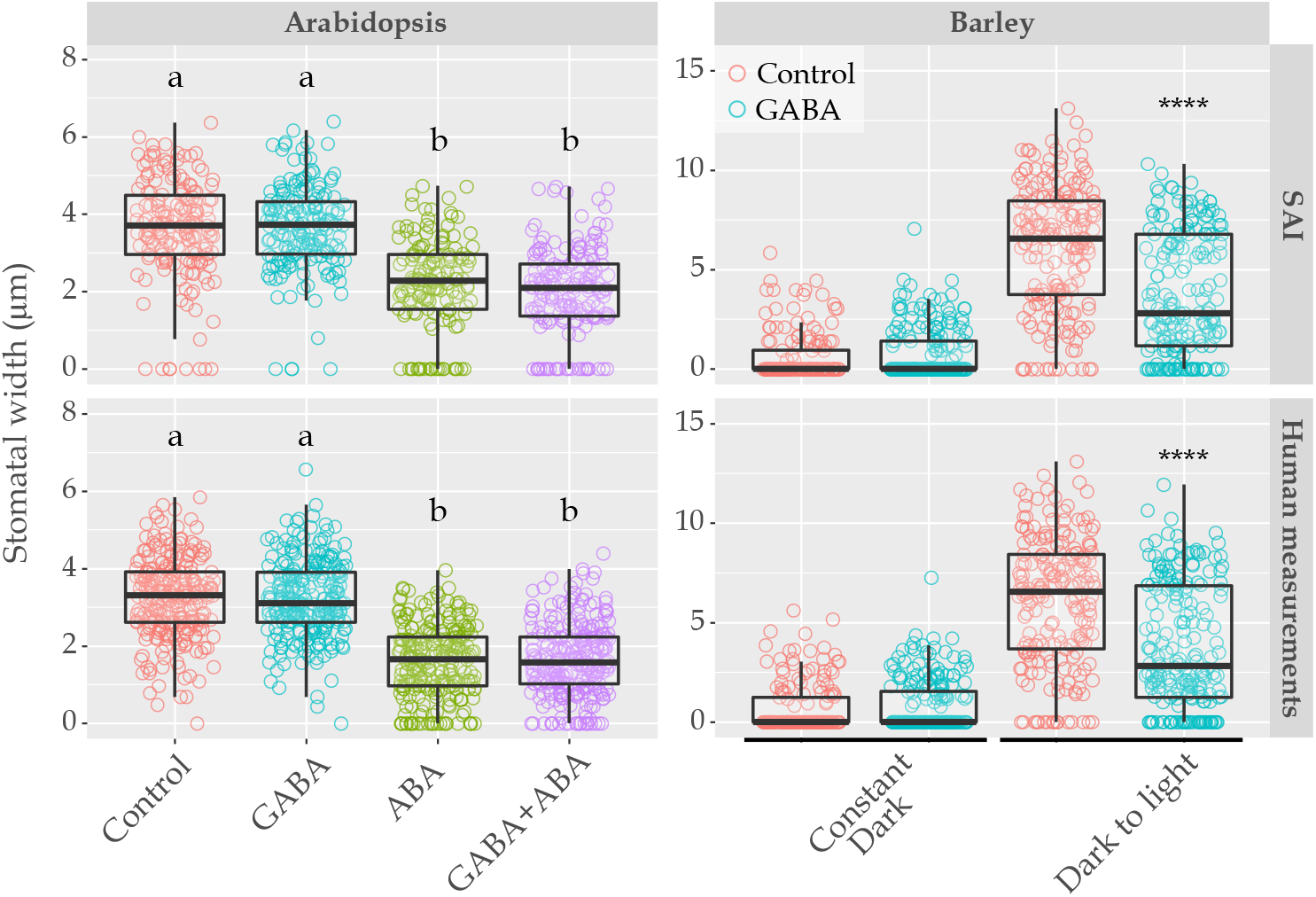
SAI predicted measurements are consistent with outcomes obtained by human researchers. Stomatal width in SAI predictions and human measurements collected with treatment under 25 μM ABA with 2 mM GABA (Arabidopsis) or 1 mM GABA (barley) during a dark-to-light transition^11^. All data was tested using one-way ANOVA followed by Tukey HSD (*N >* 140/group in Arabidopsis, *N >* 150/group in barley. a and b represent groups with no difference, *p* ≤ 0.0001 between groups, \*\*\*\**p* ≤ 0.0001).

The mean and distribution of stomatal width in each treatment group obtained using SAI were compared to the original manual measurements. SAI detected 66.37% and 91.39% of manually measured data in Arabidopsis and barley, respectively. This indicates SAI performs better when detecting barley stomata than Arabidopsis. Measurement distributions of stomatal width produced by SAI are similar in shape to those produced via manual inspection (Appendix Figure B1). Human expert 2, who was identified as the most conservative measurer, performed the human measurement of Arabidopsis stomata (Figure 3, Appendix Figure A3). Thus, the lower mean value for stomatal width obtained from human measurements was expected. When SAI measures barley samples, the distribution of measurements is almost identical to that of human measurements. This observation is likely due to the more uniform structure and higher image quality present in barley images.

### SAI significantly reduces human effort

Model inference time is predominantly limited by image resolution and computation speed. Due to this, barley data (2880×2048 in resolution) generally took a longer time to process than Arabidopsis data (2592×1944 in resolution) when using the same processor. Figure 4 shows the average time required to process a microscope image on a range of commonly available hardware.

**Fig. 4:**
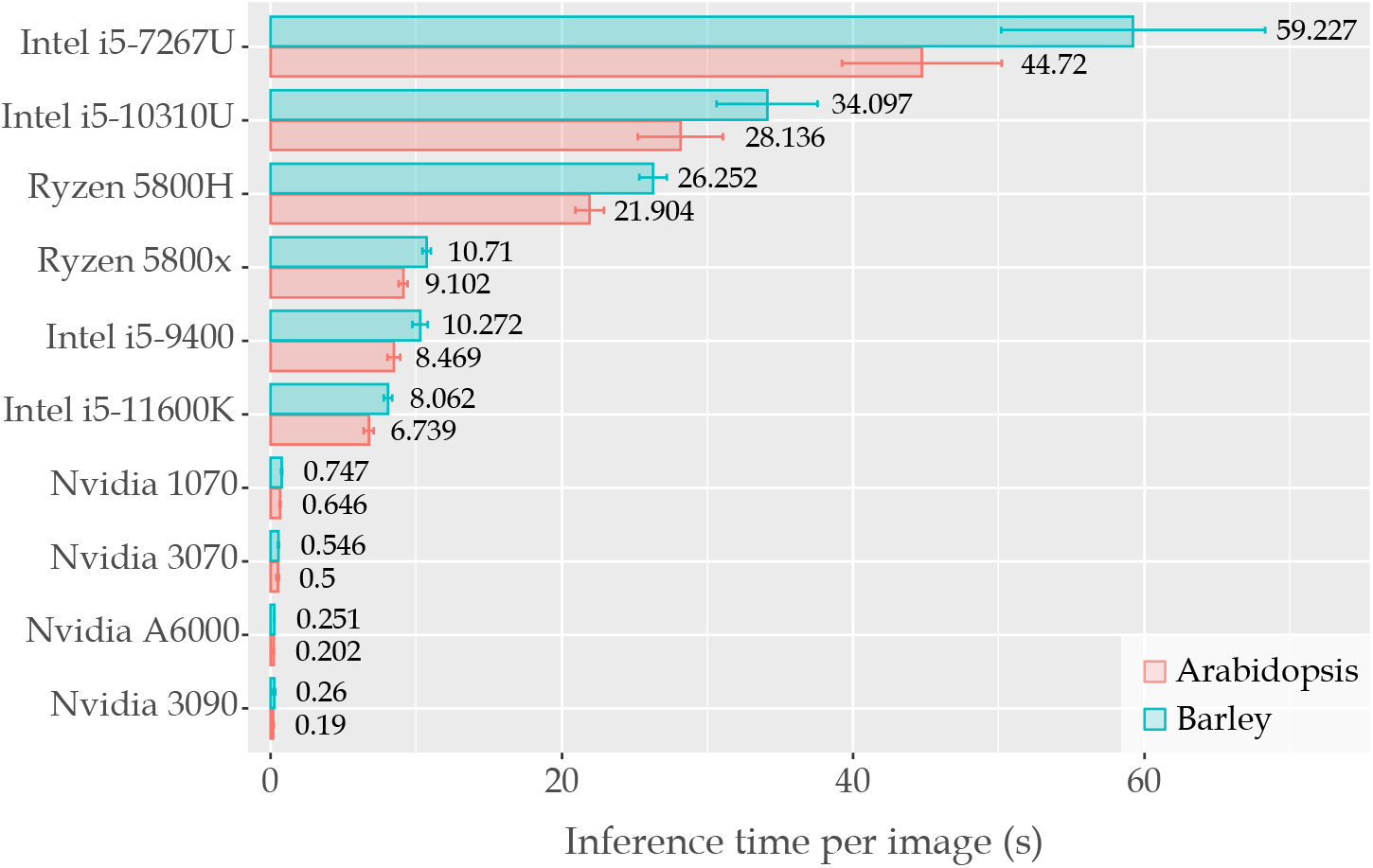
Wall clock time measured when processing a single microscope image using SAI. All processors are tested on the same image set of Arabidopsis and barley at the respective species native resolution. For all tests, the confidence threshold is set to 0.5. Mean of inference time per sample is displayed with estimated sample standard deviation. All processors were on desktop, except Intel i5-10310U, Intel i5-7267U (MacBook Pro 2017), Ryzen 5800H and Nvidia 3070 were on a laptop.

## Discussion

SAI provides a new way to analyze stomata, one of the most studied plant cell types. Our tool allows a series of stomatal features to be extracted. Specifically, opening status, complex location, width, length, and area of stomatal pores. Throughout testing, SAI generally produces better predictions on barley than Arabidopsis. This is observed for both the number of true detections made and measurement quality that reflected by CCC. This might due to image quality and leaf epidermis morphological structure. Although stomata have a relatively uniform structure, the random distribution and orientation of Arabidopsis stomata make the measurement task more challenging than for barley, which has stomata that are aligned in parallel rows with fixed orientations (Figure 1). Using the criteria outlined in Jayakody et al. (2021)^30^ to assess image quality, we found that Arabidopsis samples are rated as medium quality, whereas barley samples are considered high quality. The observed disparity in measurement quality supports the claim made in Jayakody et al. (2021)^30^ that image quality has a major impact on model performance.

The common technique used in automated stomatal pore measurement systems is to use ellipse fitting to estimate pore area, width and length from the fitted ellipse’s area, minor-axis and major-axis^22,24,27,28,32^. Ellipse fitting is limited to stomatal pores that have ovular shapes, such as those delineated by kidney-shaped guard cells (Figure 1). Plants like barley, which have stomatal pores delineated by dumbbell-shaped guard cells, do not have elliptical pores (Figure 1). Their stomatal pores resemble a coin slot, which cannot be represented accurately with an ellipse, leading to under or over estimation in derived measurements. In contrast, SAI uses direct mask segmentation of the stomata pores, which is flexible to represent any pore shape and obtain pore area by calculating masked pixel area. Moreover, the efficacy of ellipse fitting is positively correlated with the extent of stomatal pore opening^28^. Therefore, ellipse fitting cannot be used effectively for stomatal assays under experimental conditions that require measurement of stomata that are partially open or completely closed, e.g., stomata exposed to ABA, high CO_2_, H_2_O_2_ and darkness to induce stomatal closure^11,13^. SAI classified stomata before performing the measuring task and recorded the closed stomata with width and area as zero. This process includes closed stomata in the dataset and allows SAI to deal with the real-world experimental design in plant research.

From our analysis we have determined that SAI achieves *human-level* performance and when used leads to conclusions consistent with human researchers. However, SAI has many advantages compared to manual measurement. SAI produce stable and reproducible measurements. We observed that manual measurements contains rater’s bias in Average-Human/Machine Test, therefore, measurements produced by two different experts will vary (Appendix Figure A2). In contrast, SAI’s predictions are consistent, regardless of the researcher using it. This guarantees that measurements are reproducible from the same set of samples. Of particular importance to the future of plant physiology, SAI enables researchers to verify other’s conclusions without weeks of human effort. Provided that the samples from which a biologist draws their conclusions are available, SAI can produce a set of measurements within minutes. These measurements can then be used to verify claims through the application of statistical analysis.

In our experiments, SAI detects less stomata than experts. Human experts are able to use their experience to extract measurements from some stomata that are blurry, occluded or unresolved. When SAI views such samples, it will ascribe a low level of confidence to its associated measurements. To prevent false positives, where SAI predicts there is a stoma present incorrectly, we discard measurements corresponding to detections below a minimum value of confidence. Confidence is an arbitrary scale from 0 to 1 that indicates how strongly the model responded to the region. In our experience, a confidence threshold of 0.5 allows the majority of false positives to be removed while retaining valid detections. Due to this process, stomata capable of being salvaged by experts are often discarded by SAI. It is important to note that the extraction of measurements from every single stoma from an image becomes less important due to SAI’s high throughput and ability to quickly measure orders of magnitude more stomata.

Compared to manual measurement, SAI is exceptionally efficient. Depending on the confidence threshold, SAI can produce measurements from a high-resolution image in 6-12 seconds, while running on a mid-range desktop computer’s central processing unit (CPU) (Figure 4). For a human, the equivalent process takes between 2 and 5 minutes depending on image quality, the number of stomata per image, measurements required and stomatal opening status. When using a graphics processing unit (GPU), SAI further increases this disparity. On an NVIDIA GTX 1070, SAI is able to process an image every 600 milliseconds. This means that with an entry-level GPU, SAI can process hundreds of images within a minute - the equivalent of 7-9 human hours. Automatically processing hundreds of images by SAI, makes it trivial to achieve minimum measurement numbers required per treatment group for statistical testing. In fact, SAI enables researchers to increase the statistical power of their conclusions. Here, SAI decouples human effort from the number of measurements per treatment group, making measuring additional pores an attractive prospect. This enables researchers to measure previously unthinkable quantities of stomata per treatment group, allowing the law of large numbers to provide more accurate summary statistics of stomatal response.

SAI makes it possible to produce more accurate population measurements by processing a greater number of stomata in a shorter time period. This hassle-free, high resolution, time-efficient data acquisition assistant has the potential to accelerate research that has a major impact on plant physiology. Moreover, SAI’s ability to learn how to measure stoma in both barley and Arabidopsis gives us confidence in its ability to do so in other species. Towards this, we provide an additional model with SAI which has been trained on a combined species data set. As pores share some common visual features, this model can be used as a starting point for researchers wishing to use SAI on a new type of plant. To do this, a set of measured examples that conform to our annotation format could be used to *fine tune* the provided combined species model. In this study, we only consider SAI’s use in measuring pore features. However, given sufficient labelled data, extension to measurement of other relevant cell structures would be possible. More generally, SAI could be used to measure other structures captured via microscopy.

SAI is a reliable, fast and simple solution to automate stomatal measuring for plant biologists via a user-friendly web app (online demo is available at https://sai.aiml.team, full version is available at https://github.com/XDynames/SAI-app). SAI is a new tool that can free researchers from labour intensive low-throughput measuring tasks; accelerating the speed of physiology-based research, regardless of the shape of the stomatal pore.

## Methods

### Data annotation and Modeling

*Arabidopsis thaliana* ecotype Col-0 and barley (*Hordeum vulgare*, Barke) were prepared as plant material^11^. Arabidopsis images were captured using Axiophot Pol Photomicroscope (Carl Zeiss). A Nikon DS-Fi3 digital camera with a Nikon diaphot 200 inverted microscope was used to capture barley images. All images were annotated using RectLabel (version 3.03.8, https://rectlabel.com). Creation of pore feature annotations followed the procedure of manual measurement carried out in Fiji-ImageJ. In a given annotation, two labels were ascribed to each stoma (Figure 5). A bounding box which contains a single stoma with associated opening status (open or closed). Measurements of the stomatal pore were also taken. These were recorded as a polygon or a line for open and closed stomata respectively. All information was organised for compatibility with Microsoft Common Objects in Context (MS-COCO); which is a widely used computer vision benchmark^34^. Summary statistics of the created database of microscopy images are presented in Table 1.

**Table 1:** Summary metrics for stomatal pore dataset used in model training and evaluation.

|  | Slide Images |  | Stoma Instances |  |
| --- | --- | --- | --- | --- |
|  | Train | Validation | Train | Validation |
| Arabidopsis | 200 | 42 | Open: 974<br>Closed: 293 | Open: 235<br>Closed: 55 |
| Barley | 150 | 33 | Open: 1000<br>Closed: 692 | Open: 268<br>Closed: 89 |

**Fig. 5:**
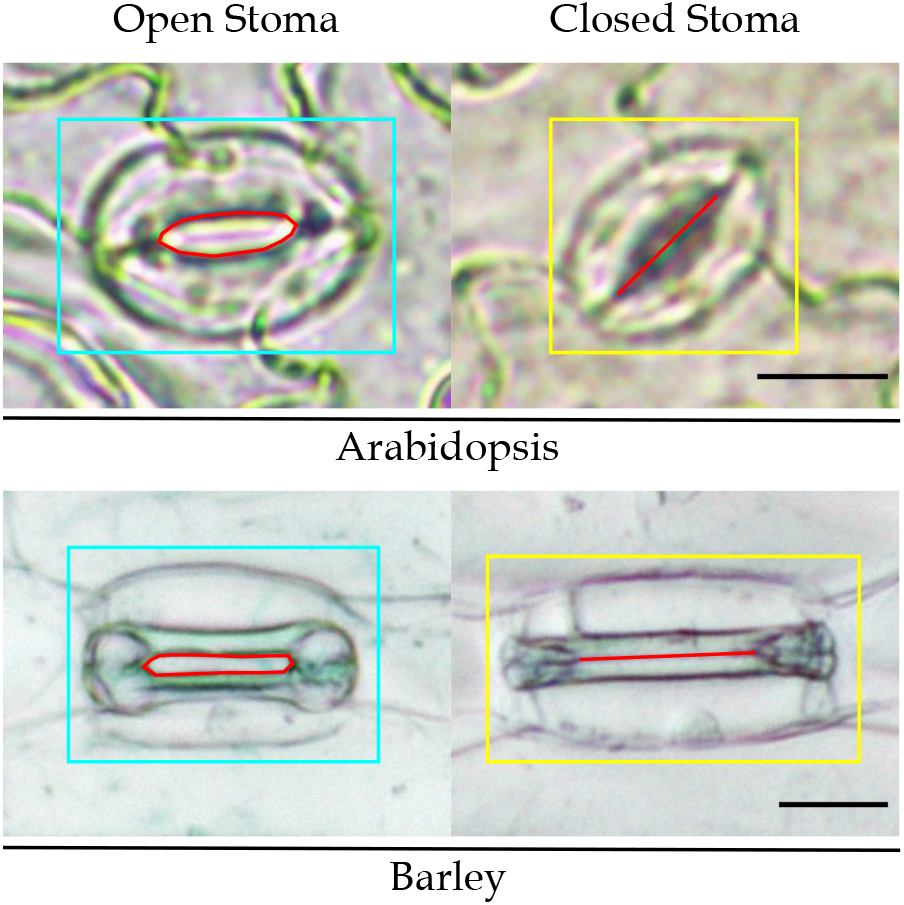
Annotation examples of Arabidopsis and barley stomata. Bounding boxes contain a single Arabidopsis or barley stoma (i.e. a pair of guard cells and a pair of subsidiary cells if from a relevant plant) and its opening status is determined in different label (open stoma in cyan, closed stoma in yellow). The red polygon and line present defined stomatal pore in each annotation. Scale bar represents 10 μm in Arabidopsis and 20 μm in barley.

Traditionally, researchers inspect each captured microscope image and measure relevant structures of a stoma. The measurement procedure will depend on the pore opening status. The area of an open stoma is measured by drawing a polygon that encloses its *mouth*. To determine the pore’s width and length, researchers either directly measure or apply fitting methods to the aforementioned polygon. Closed pore lengths are acquired by selecting points which mark the beginning and end of their tightly shut *mouth*. For a computer model to emulate a researcher performing the measurement, it must: localise target structures within a sample; comment on their state and gather relevant measurements. To enable this, we have reformulated each one of these tasks into a computer vision task. A researcher’s initial identification of stomata and their opening status is re-framed as object detection. Object detection consists of drawing boxes around salient objects and predicting the enclosed object’s semantics. Drawing polygons indicating stomatal openings maps to segmentation, which highlights regions of interest within images. Selecting end points for a stomatal pore is analogous to keypoint detection, which reduces visual features of interest to a defining pixel. Each of these tasks has a library of possible models capable of solving them individually. By requiring an all-inclusive solution, candidates are significantly reduced. Mask R-CNN represents an incremental change atop of an already established series of deep-learning architectures^35–38^. This iteration comes armed with requisite predictive powers for our physiological needs. Through the use of specialized predictive *heads*, Mask-RCNN is capable of learning object detection, segmentation and keypoint detection in tandem.

Deep-learning models were built using Detectron 2; an open-source framework sitting on top of Pytorch. Both of these packages were created by Facebook’s Artificial Intelligence Research division (FAIR)^39,40^. Adaptions were made to FAIR’s Mask R-CNN model to better suit stomatal measurement. Specifically, increasing the resolution of prediction heads responsible for segmentation and keypoint detection. Mean average precision (*mAP*), as defined in the MS-COCO challenge, was used to evaluate and compare models on all tasks^34^. Justification and verification of model design choices and training regimes are presented in Appendices C-J.

### Average-Human/Machine Test

To determine whether SAI predictions were consistent with human measurement, an Average-Human/Machine Test was designed. In total, 35 microscopy images, 15 of barley and 20 of Arabidopsis, were collated as a test dataset (summarized in Table 2). Four plant stomata morphology experts participated by manually measuring the 35 random selected images. The mean measurements of 4 human experts on each stoma were used as a *human-level* reference. This reference was used to quantify the extent to which a single researcher may vary within their own judgement and assess SAI. Participants used the same annotation schema outlined above data preparation. To understand how measurements retrieved from images change with respect to their measurers, all measurements were matched to a *human-level* reference at the single stoma level. Differences between measurement sources against the *human-level* reference were visualized with scatter plots and quantified by relative error. To evaluate the agreement between SAI/human experts and *human-level* reference, the concordance correlation coefficient (CCC; ranging from −1 to 1) was applied^33^. Furthermore, the means of each measurement, for each measurer, was compared using one-way ANOVA with Tukey HSD.

**Table 2:** Summary of Average-Human/Machine Test dataset.

|  | Images | Stomata |  |
| --- | --- | --- | --- |
|  |  | Open | Closed |
| Arabidopsis | 20 | 129 | 20 |
| Barley | 15 | 109 | 66 |

### SAI in practice

To support our claim that SAI is a replacement for traditional measurement methods, we demonstrate that scientific conclusions drawn from measurements produced by SAI align with those of expert physiologists. Manually measured image datasets of both Arabidopsis and barley were obtained from Xu et al. (2021)^11^. Two different experimental designs were selected to evaluate SAI’s real-world performance: the 25 μM ABA with presence and absent of 2 mM GABA for Arabidopsis and dark-to-light transition with or without 1 mM GABA for barley. Reference measurements for the two datasets were made by different researchers. Arabidopsis measurements were taken by human expert 2. Barley samples were measured by human expert 4. Measurements produced by SAI were subjected to the same statistical tests used in Xu et al. (2021)^11^ to examine whether SAI enables consistent conclusions as those reached by expert measurements.

### Inference time assay

The efficiency of SAI compared to manual labelling was tested on a range of commonly available computer hardware. We do this using the same set of sample images used in the Average-Human/Machine Test (Table 2). Time to processes each image was recorded and used to estimate the average inference time and sample standard deviation for each processor. These measures were then used to compare their throughput.

