## Supplementary Data for "SAI: Fast and automated quantification of stomatal parameters on microscope images"

Part of this work was done when Hao Chen and Chunhua Shen were with University of Adelaide.

[#] Equal contribution.

SUPPLEMENTARY INFORMATION

### Appendix A Supplementary Data of Average-Human/Machine Test

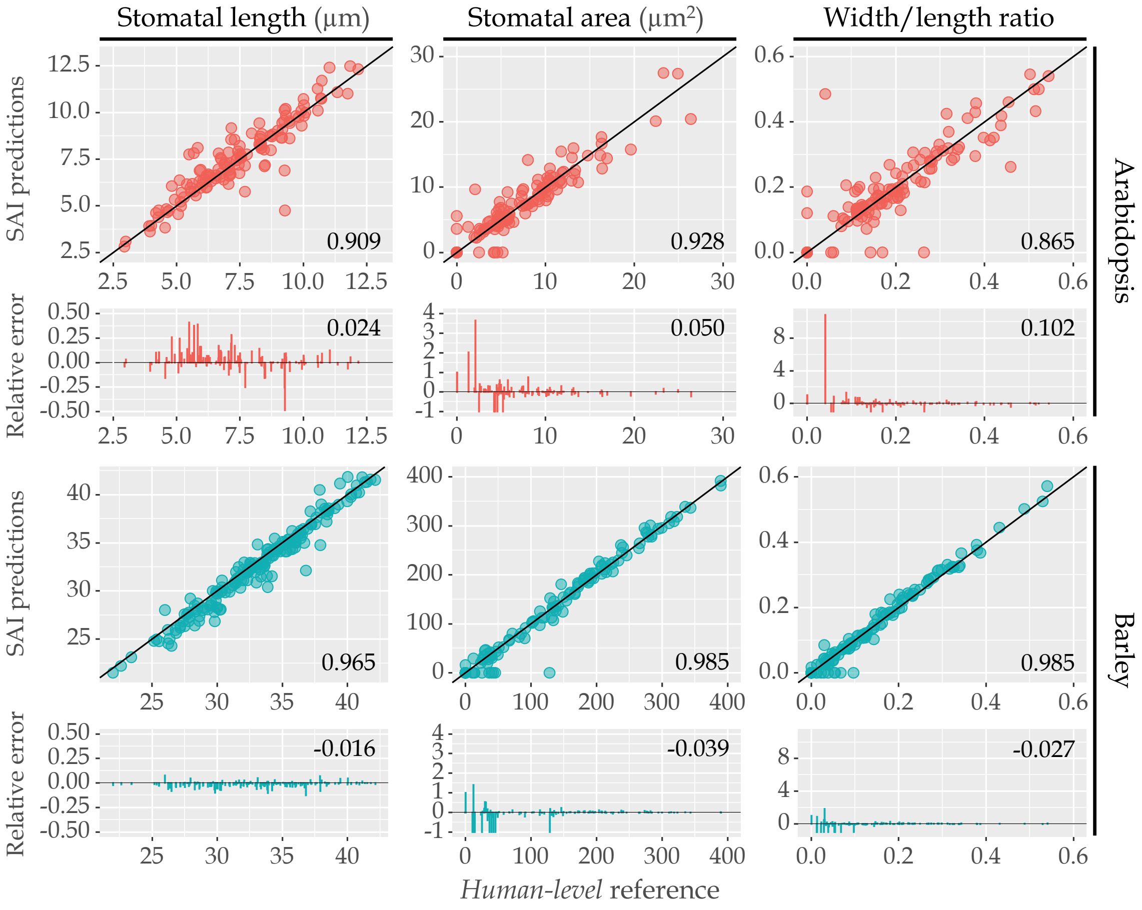


**Fig. A1: SAI prediction vs human-level reference set for Arabidopsis and barley stomatal length (µm), area (µm$^2$) and width/length ratio.** Measurements from 4 human experts on stomata morphology were collected and average length and area of each stoma were calculated as the human-level reference. SAI is compared against the reference and the concordance correlation coefficient (ranging from $-1$ to $1$) were calculated as the determination of the accuracy performance. The corresponding relative error (RE) to human-level reference was displayed on the corresponding sub-figure with mean RE calculated. Data points are colour coded according to annotation label. (Arabidopsis: $N > 120$, barley: $N > 160$)

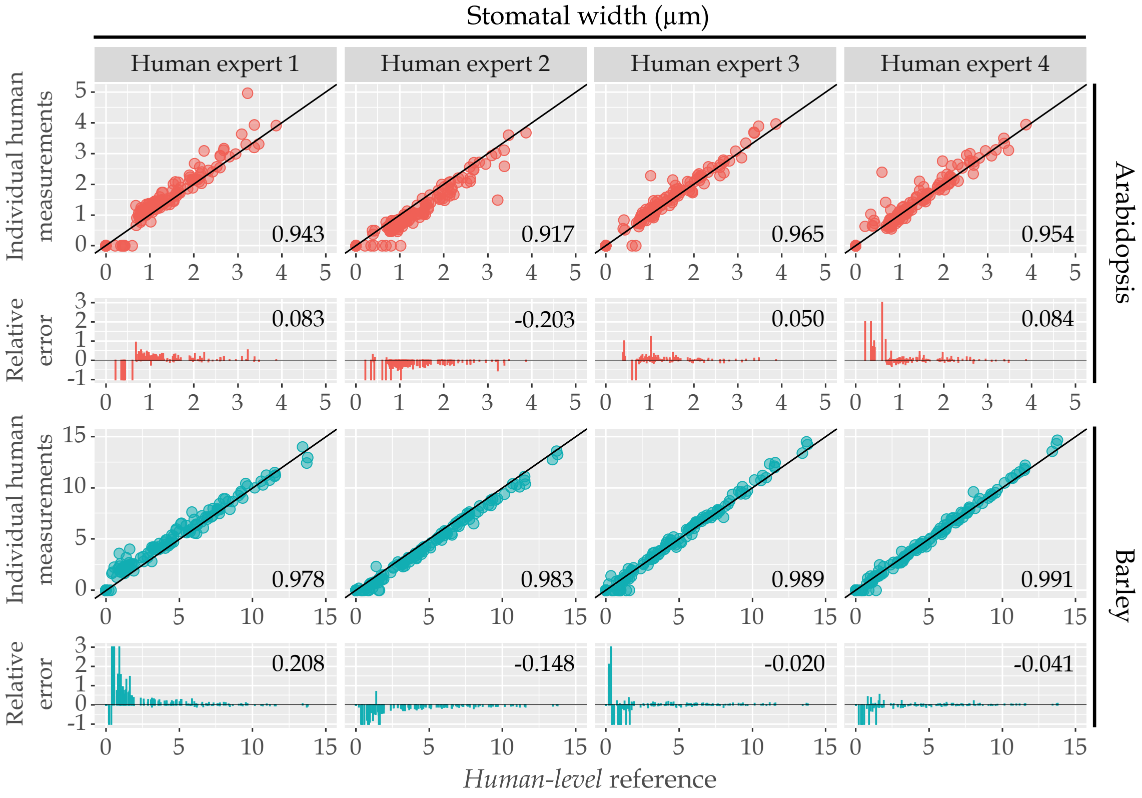
Stomatal width (µm)
Human expert 1
Human expert 2
Human expert 3
Human expert 4
Individual human measurements
Relative error
Arabidopsis
Barley
0.943
0.917
0.965
0.954
0.083
-0.203
0.050
0.084
0.978
0.983
0.989
0.991
0.208
-0.148
-0.020
-0.041
Human-level reference

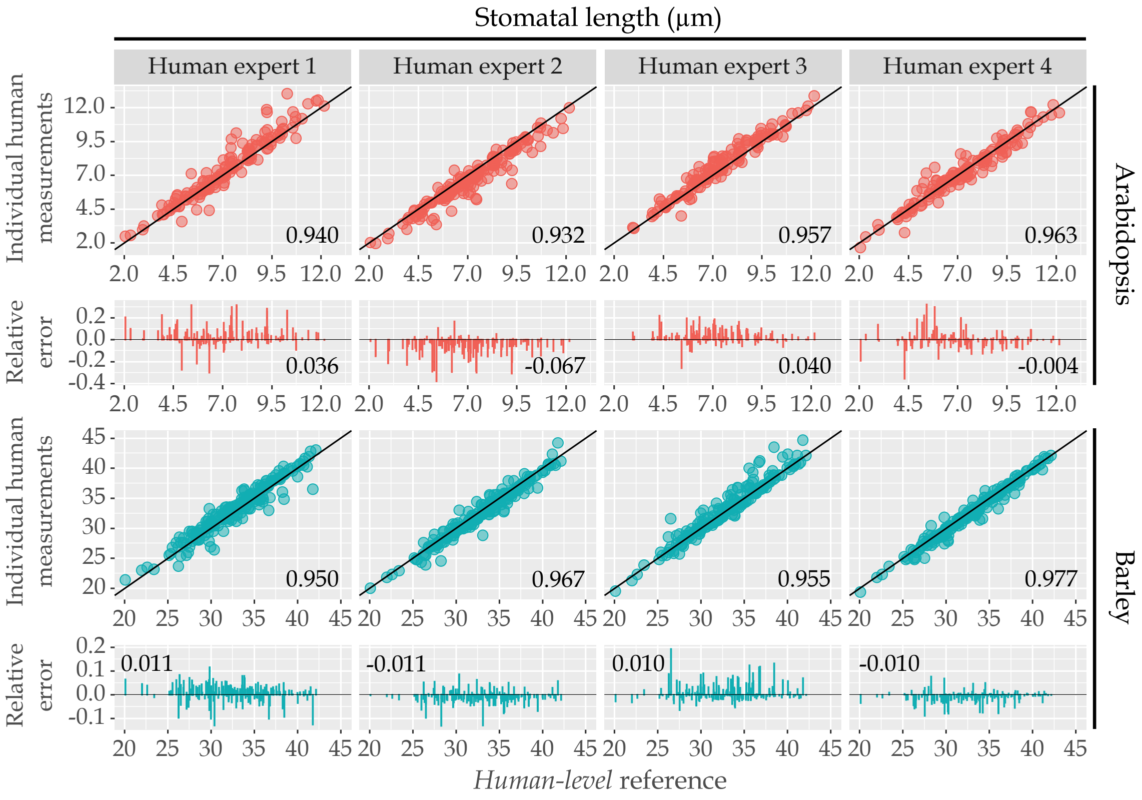
Stomatal length (µm)
Human expert 1
Human expert 2
Human expert 3
Human expert 4
Individual human measurements
Relative error
Arabidopsis
Barley
0.940
0.932
0.957
0.963
0.036
-0.067
0.040
-0.004
0.950
0.967
0.955
0.977
0.011
-0.011
0.010
-0.010
Human-level reference

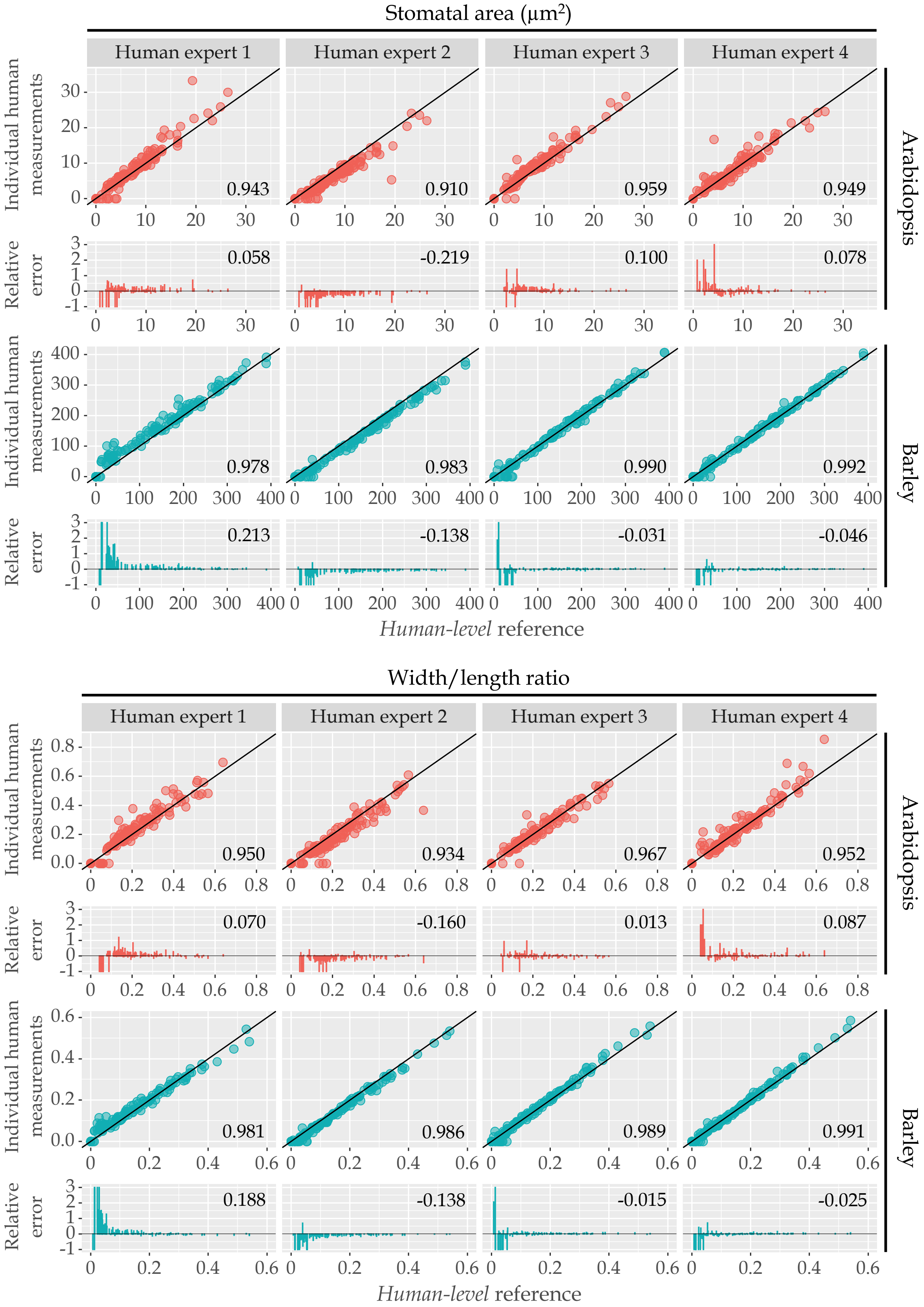


**Fig. A2: Individual human measurements vs *human-level* reference set for Arabidopsis and barley with corresponding relative error of stomatal width, length, area and width/length ratio.** Measurements from 4 human experts on stomata morphology are compare against the *human-level* reference (the average of human measurements) in each stoma and the concordance correlation coefficient (ranging from $-1$ to 1) were calculated as the determination of the accuracy performance. Stomatal width, length, area and width/length ratio from all experts were matched with human reference to calculate relative error. Mean of relative error were calculated and displayed on corresponding sub-figure. (Arabidopsis: $N > 120$, barley: $N > 160$)

Table A1: Summary of measured stomata in number and mean value of corresponding measuring feature.

| Species | Predictor | $N$ | Mean | | | |
|---|---|---|---|---|---|---|
| | | | length | width | area | width/length ratio |
| Arabidopsis | Human expert 1 | 144 | 7.489 | 1.411 | 7.643 | 0.213 |
| | Human expert 2 | 149 | 6.770 | 1.028 | 5.547 | 0.169 |
| | Human expert 3 | 132 | 7.719 | 1.361 | 8.102 | 0.194 |
| | Human expert 4 | 139 | 7.225 | 1.322 | 7.200 | 0.209 |
| | SAI | 127 | 7.531 | 1.312 | 7.293 | 0.192 |
| Barley | Human expert 1 | 175 | 33.232 | 3.657 | 112.002 | 0.112 |
| | Human expert 2 | 174 | 32.489 | 2.929 | 88.027 | 0.094 |
| | Human expert 3 | 172 | 33.203 | 3.416 | 101.689 | 0.109 |
| | Human expert 4 | 174 | 32.579 | 3.35 | 100.300 | 0.107 |
| | SAI | 166 | 32.578 | 3.302 | 100.155 | 0.106 |

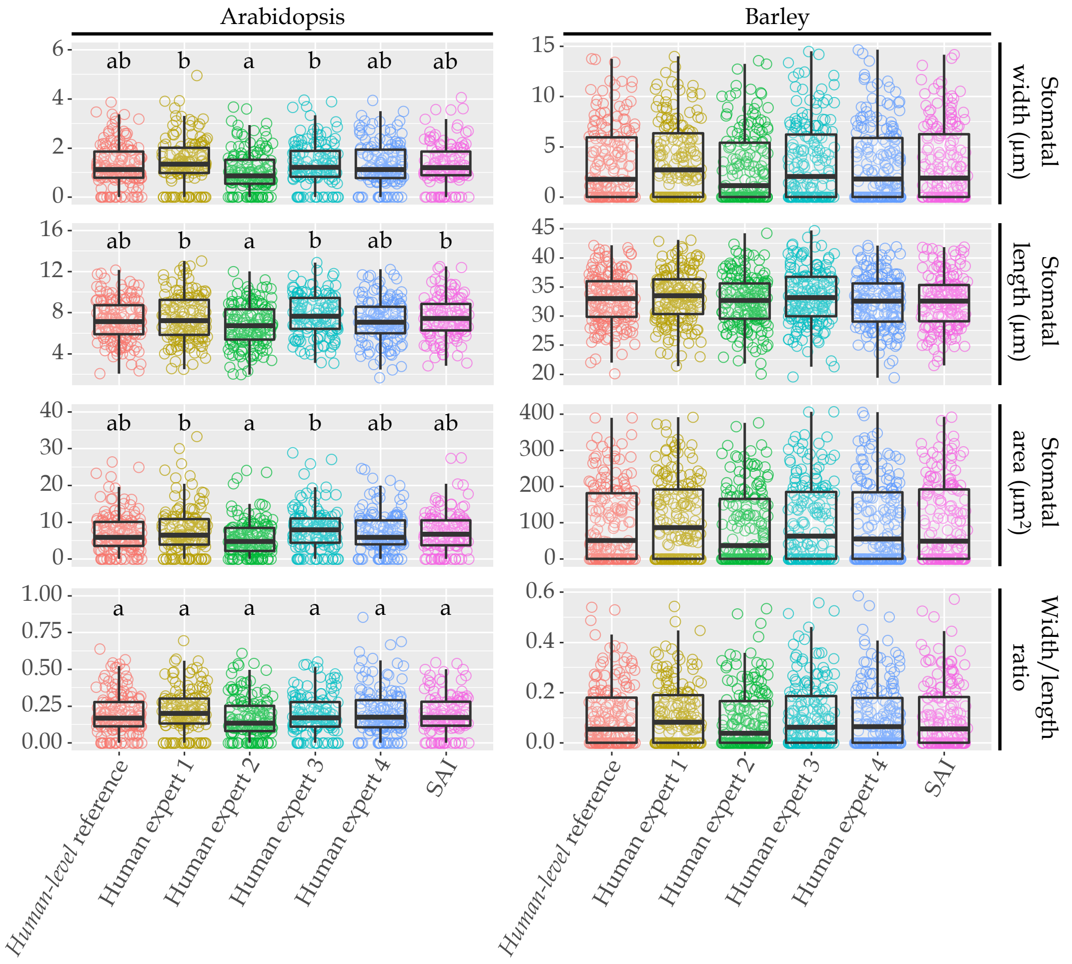


**Fig. A3: Measurement comparison of stomatal width, length, area and width/length ratio in Arabidopsis and barley.** Individual stomatal measurement and median visualised in box plot from *human-level* reference (the average of human measurements), 4 human experts on stomata morphology and SAI presented with one-way ANOVA with Tukey HSD Test. No differences found between source of measurements in barley, a and b represent groups without significant difference in Arabidopsis, $p \leq 0.05$ between group.

### Appendix B Supplementary Data of Human Processed Datasets

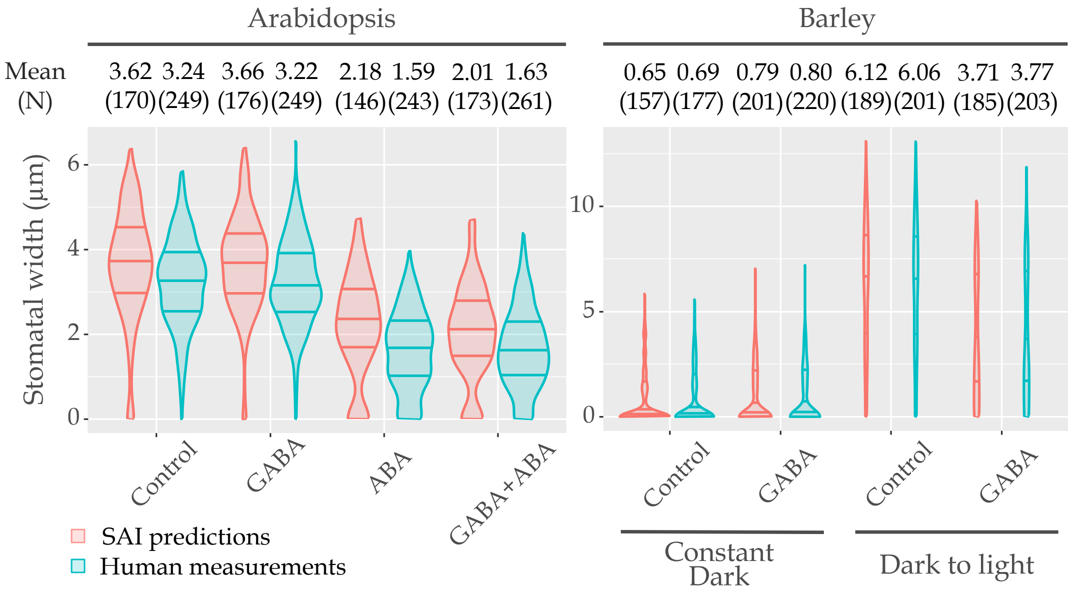


**Fig. B1: Mean of stomatal width and measurement distribution illustration.** The violin plot presents the distribution shape of measurements; and the interquartile range with median are marked as the line. The mean of stomatal width N were calculated in corresponding group.

### Appendix C Default Training Procedure

Models are trained for 90,000 iterations each. Learning rate is started at a value of 0.0006 ramping up linearly to 0.005 over the first 1,000 steps. This is then decreased by a factor of ten at 15,000 and 25,000 steps. Random cropping and horizontal flipping were employed as augmentation; adding additional variation to examples. Standard input and evaluation resolution used is $800 \times 1333$.

### Appendix D Model performance

Stomatal measurement was carried out on two plant species: Arabidopsis and barley. Validation samples of barley stoma were kept at their native resolution of $2880 \times 2048$ when used to evaluate model measurements. Our model obtained a mean average precision ($\mathrm{mAP}$) of $84.91\% \pm 0.59$ for bounding boxes, $70.44\% \pm 0.12$ for segmentation masks and $84.28\% \pm 0.17$, $67.09\% \pm 10.60$ for keypoint detections on open and closed samples respectively. Arabidopsis measurements were evaluated on validation images at their native resolution of $2592 \times 1944$. Models trained to measure Arabidopsis stomatal pores obtained a $\mathrm{mAP}$ of $73.81\% \pm 1.14$ for bounding boxes, $44.18\% \pm 0.23$ for segmentation masks and $53.11\% \pm 1.53$, $32.21\% \pm 4.5$ for keypoint detections on open and closed samples respectively. Mean and sample standard deviation estimates were obtained by training five models with different random initialization. Reported scores are for raw predictions without post-processing techniques introduced as part of using the model within the measurement application.

### Appendix E Batch Size

A series of models were trained to explore the effect of input batch size on predictive power. Each model underwent the same training regime, outlined in Appendix C, but with a modified batch size. Evidenced by Table E1, the number of input images during training has no impact on the learned model's inference. Therefore, if another user wishes to train this architecture, they need only have hardware capable of processing a single images at a time.

Table E1: Model prediction scores when training with the default strategy outlined in Appendix C for various input batch sizes.

| Batch Size | Bounding Box mAP % | Segmentation mAP % | Keypoints mAP %(Open, Closed) |
|---|---|---|---|
| 1 | 82.34 | 60.92 | 71.14 (79.43, 62.86) |
| 2 | 82.49 | 58.30 | 71.00 (78.55, 63.45) |
| 4 | 81.30 | 58.59 | 70.59 (77.91, 63.27) |
| 6 | 81.45 | 58.16 | 70.46 (78.46, 82.46) |
| 8 | 80.94 | 59.60 | 71.75 (79.32, 64.19) |
| 12 | 80.00 | 57.92 | 70.37 (79.12, 61.62) |

### Appendix F Keypoint Head Complexity

Default configuration of Mask-RCNN's keypoint detection head is tailored towards human pose estimation or identifying facial keypoints. In this case, models are required to localise multiple points, either across a person or on their face. For our use case, only two points need to be localised. Due to this, it is argued that a reduction of complexity will not severely impact model performance on stomatal pores. Results of experiments to explore this are presented in Table F1. Deeper and wider are said to be more complex than shallow and narrow. In Table F1 we can see our suspicion, that keypoint head complexity can be reduced without significant impact, is supported. Bounding box and segmentation tasks are not impacted either; as should be expected. Keypoint localisation shows a slight decrease in accuracy when moving from a convolution width of 512 to 256 but only for a layer depth of 8. In all other cases, differences in ability are within $\pm 1.5\%$. Motivated by these results,

Table F1: Comparison of limiting keypoint head depth and width.

| Depth | Width | Bounding Box mAP % | Segmentation mAP % | Keypoints mAP % (Open, Closed) |
|---|---|---|---|---|
| 2 | 256 | 80.17 | 56.38 | 70.01(79.42, 60.59) |
| | 512 | 79.29 | 57.25 | 69.61(77.64, 61.59) |
| 4 | 256 | 80.12 | 57.75 | 71.37(78.34, 64.41) |
| | 512 | 80.19 | 56.73 | 70.53(78.53, 62.53) |
| 8 | 256 | 79.97 | 56.6 | 70.44(78.09, 62.78) |
| | 512 | 81.68 | 57.74 | 73.52(80.28, 66.77) |

Note: Pooler, training and testing image resolutions are fixed at $14 \times 14$, $320 \times 800$ and $800 \times 1333$ respectively. All models use the default training strategy outlined in Appendix C, on the barley pore dataset.

a reduced head, with depth of 2 and width of 256, is used. This decision reduces video memory constraints for further modifications.

### Appendix G Keypoint and Mask Head Pooler Resolution

In most cases, crops containing pores generated by the region proposal network are of high resolution in comparison to poolers used by default in Mask-RCNN. A region proposal large enough to contain a cell will have dimensions of $300 \times 200$ pixels; default poolers reduce these proposals down to $14 \times 14$ for further processing. This means that prediction heads will need to make decisions about a much larger region from this considerably compressed summary. Ideally, predictions would be made using a feature map with full proposal resolution, but two barriers prevent this: memory consumption and uniform matrix size. To understand the benefits of increasing pooler output size, a series of models were trained with different keypoint and mask head pooler resolutions, shown in Tables G1 & G2. A modified pooler that enables non-square dimensions is explored as an option in keypoint heads to better reflect closed stomatal pore crop's rectangular aspect. Evidenced by Table G1 increasing resolution from $14 \times 14$ shows immediate benefit, however further square increase tends to only benefit prediction on closed stomatal pores. The same benefit as doubling both height and width can be achieved by using a rectangular pooler, where only one dimension is increased. From example, instead of increasing from $28 \times 28$ to $56 \times 56$, changing to $28 \times 56$ benefits a model's ability to predict on closed stomatal pores more without impacting predictions made on open examples.

Table G1: Trails highlighting the impact of keypoint head pooler resolution on model performance.

| Pooler Resolution | Bounding Box mAP % | Segmentation mAP % | Keypoints mAP %(Open, Closed) |
|---|---|---|---|
| $14 \times 14$ | 80.17 | 56.38 | 70.01 (79.42, 60.59) |
| $28 \times 28$ | 82.70 | 58.92 | 72.72 (81.37, 63.96) |
| $56 \times 56$ | 82.41 | 58.78 | 73.73 (81.11, 66.35) |
| $112 \times 112$ | 82.40 | 59.26 | 73.12 (79.68, 64.90) |
| $14 \times 28$ | 82.04 | 57.82 | 72.29 (79.68, 64.90) |
| $28 \times 56$ | 79.68 | 59.27 | 73.21 (81.37, 65.04) |
| $56 \times 112$ | 81.07 | 58.84 | 75.33 (80.47, 70.18) |

Note: All models are trained using default training procedure outlined in Appendix C. Batch size varies across trails but has no significant impact on a model's predictive power, see Appendix E.

Table G2: Experiments isolating mask head pooler resolution's effect on task performance.

| Pooler Resolution | Bounding Box mAP % | Segmentation mAP % | Keypoints mAP %(Open, Closed) |
|---|---|---|---|
| $14 \times 14$ | 79.68 | 59.27 | 73.21 (81.37, 65.04) |
| $28 \times 28$ | 81.42 | 67.84 | 75.25 (81.84, 68.67) |
| $56 \times 56$ | 82.22 | 70.03 | 73.71 (81.51, 65.91) |

Note: All models are trained using default training procedure outlined in Appendix C. A batch size of 2 is used, and all models use a $2 \times 256$ keypoint head with pooler resolution of $26 \times 56$.

As masking stomatal openings is required only when a pore is open, square pooler resolutions are considered for mask heads. Table G2 shows that an initial benefit to segmentation performance is seen in response to the pooler resolution, but as higher values are used diminishing returns are observed.

### Appendix H Image Resolution

By using a smaller input resolution, the time for a model to train can be reduced. However, when a model is taught with examples at low resolution, it may struggle to interpret patterns that are scaled up in a high resolution example of the same image. Ideally, inference is performed at the native capture resolution of the microscopy setup. This eliminates scaling artefacts which introduce noise to samples. In Table H1 the impact of training time resolution on inference metrics is explored. The aforementioned effect of training on small and inferring on high resolutions is observed for both pooler resolutions tested. As training resolution is increased, test time performance improves sharply and then begins to show diminishing returns. Providing a low value for the minimum crop size with an almost native maximum tends to show either no benefit or slight deficit in performance. From these trails $1200, 2048$ is identified as a strong candidate as a default crop size for training when inference resolution is close to capture resolution.

Table H1: Quantifying the impact of training crop resolution on final model predictive power.

| Pooler Resolution | Training Crop Size | Bounding Box mAP % | Segmentation mAP % | Keypoints mAP %(Open, Closed) |
|---|---|---|---|---|
| | 320, 800 | 54.24 | 25.71 | 39.53 (46.32, 32.74) |
| | 800, 1200 | 83.41 | 56.54 | 75.44 (81.43, 69.45) |
| $14 \times 14$ | 1200, 2048 | 84.42 | 61.10 | 78.92 (84.66, 73.17) |
| | 320, 2048 | 84.93 | 59.19 | 77.98 (85.54, 70.43) |
| | 320, 3000 | 84.83 | 59.85 | 77.94 (84.55, 71.33) |
| | 320, 800 | 25.96 | 30.15 | 28.16 (10.36, 45.96) |
| | 800, 1200 | 79.94 | 38.17 | 62.20 (73.55, 50.85) |
| $28 \times 56$ | 1200, 2048 | 83.70 | 58.69 | 77.21 (83.37, 71.17) |
| | 320, 2048 | 83.84 | 58.40 | 77.60 (83.76, 71.53) |
| | 320, 3000 | 84.16 | 60.11 | 76.40 (83.26, 69.54) |

Note: Models with pooler resolution of $14 \times 14$ use keypoints heads with high complexity prediction heads, $8 \times 512$ convolutions. $28 \times 56$ used minimal complexity heads with $2 \times 256$ convolution layers. Training crop size is provided to show the minimum and maximum size of crops taken from the image that are then re-scaled to a consistent size prior to the model seeing them. All models are trained using default training procedure outlined in C and evaluated at native resolution on the validation set.

### Appendix I Keypoint Head Complexity at Higher Resolution

Previously, it was shown that a reduction in head complexity had very little impact on final model predictive power. This conclusion is re-examined under increased training resolution. Experiments done in Appendix F are repeated using $1200 - 2048$ training crop sizes and evaluated at native resolution input. It is observed in Table I1 that without significant change to mAP on open samples, closed stomata predictions can be boosted by $\approx 8\%$ mAP with increased training crop size.

Table I1: Comparison of keypoint head depth and width with increased pooler and input resolutions.

| Depth | Width | Bounding Box mAP % | Segmentation mAP % | Keypoints mAP % (Open, Closed) |
|---|---|---|---|---|
| 2 | 256 | 85.37 | 70.51 | 73.06 (84.55, 61.57) |
| | 512 | 85.15 | 71.4 | 73.9 (84.74, 63.06) |
| 4 | 256 | 84.67 | 70.58 | 73.99 (84.48, 63.52) |
| | 512 | 85.15 | 71.4 | 73.9 (84.76, 63.06) |
| 6 | 256 | 85.47 | 70.47 | 76.18 (84.52, 67.85) |
| | 512 | - | - | - |
| 8 | 256 | 84.94 | 70.15 | 76.44(83.68, 69.21) |
| | 512 | - | - | - |

Note: All models were trained using the final strategy outlined in Appendix J, on the barley pore dataset. Omitted entries exceeded 32GB VRAM and therefore could not be evaluated.

### Appendix J Final Schema

Based on the above ablation studies, the model used in comparisons to human annotation performance has a keypoint and mask pooler resolution of $56 \times 112$ and $56 \times 56$ respectively. The keypoint head's width is reduced to $256$ but a depth of $8$ is retained. Training random crop short edge sizes varied from $1200$ to $2048$ and an input batch size of two was used. Length of the learning rate warm up was increased from $1000$ to $7500$ to mitigate early divergence. All other learning rate, scheduling, step number and augmentation strategies remain as outlined in Appendix C. When training on Arabidopsis samples, it was found to be beneficial to include a larger difference between the minimum and maximum crop sizes while training. These models were exposed to image crops as small as 320 and as large as their native size of 1944 pixels. The large variation present in keypoint detection on closed pores indicates any further efforts should be targeted toward this task.
